# Afterload-induced Decreases in Fatty Acid Oxidation Develop Independently of Increased Glucose Utilization

**DOI:** 10.1101/2024.09.17.613531

**Authors:** Hande C. Piristine, Herman I. May, Nan Jiang, Daniel Daou, Francisco Olivares-Silva, Abdallah Elnwasany, Pamela Szweda, Luke Szweda, Caroline Kinter, Michael Kinter, Gaurav Sharma, Xiaodong Wen, Craig R. Malloy, Michael E. Jessen, Thomas G. Gillette, Joseph A. Hill

**Affiliations:** Department of Internal Medicine (Cardiology), Harry S. Moss Heart Center, UT Southwestern Medical Center, Dallas, TX, USA; Department of Molecular Biology, UT Southwestern Medical Center, Dallas, TX, USA; Simmons Comprehensive Cancer Center, UT Southwestern Medical Center, Dallas TX, USA; Aging and Metabolism Research Program, Oklahoma Medical Research Foundation, Oklahoma City, OK; Department of Cardiovascular and Thoracic Surgery, UT Southwestern Medical Center, Dallas, TX, USA; Advanced Imaging Research Center, UT Southwestern Medical Center, Dallas, TX, USA; Department of Internal Medicine, UT Southwestern Medical Center, Dallas, TX, USA; VA North Texas Health Care System, Dallas, TX, USA

**Keywords:** cardiac hypertrophy, mitochondrial metabolism, metabolic flexibility, heart failure, pyruvate dehydrogenase complex

## Abstract

**Background:** Metabolic substrate utilization in HFpEF (heart failure with preserved ejection fraction), the leading cause of heart failure worldwide, is pivotal to syndrome pathogenesis and yet remains ill defined. Under resting conditions, oxidation of free fatty acids (FFA) is the predominant energy source of the heart, supporting its unremitting contractile activity. In the context of disease-related stress, however, a shift toward greater reliance on glucose occurs. In the setting of obesity or diabetes, major contributors to HFpEF pathophysiology, the shift in metabolic substrate use toward glucose is impaired, sometimes attributed to the lower oxygen requirement of glucose oxidation versus fat metabolism. This notion, however, has never been tested conclusively. Furthermore, whereas oxygen demand increases in the setting of increased afterload, myocardial oxygen availability remains adequate for fatty acid oxidation (FAO). Therefore, a “preference” for glucose has been proposed.

**Methods and Results:** Pyruvate dehydrogenase complex (PDC) is the rate-limiting enzyme linking glycolysis to the TCA cycle. As PDK4 (PDC kinase 4) is up-regulated in HFpEF, we over-expressed PDK4 in cardiomyocytes, ensuring that PDC is phosphorylated and thereby inhibited. This leads to diminished use of pyruvate as energy substrate, mimicking the decline in glucose oxidation in HFpEF. Importantly, distinct from HFpEF-associated obesity, this model positioned us to abrogate the load-induced shift to glucose utilization in the absence of systemic high fat conditions. As expected, PDK4 transgenic mice manifested normal cardiac performance at baseline. However, they manifested a rapid and severe decline in contractile performance when challenged with modest increases in afterload triggered either by L-NAME or surgical transverse aortic constriction (TAC). This decline in function was not accompanied by an exacerbation of the myocardial hypertrophic growth response. Surprisingly, metabolic flux analysis revealed that, after TAC, fractional FAO decreased, even when glucose/pyruvate utilization was clamped at very low levels. Additionally, proteins involved in the transport and oxidation of FFA were paradoxically downregulated after TAC regardless of genotype.

**Conclusions:** These data demonstrate that cardiomyocytes in a setting in which glucose utilization is robustly diminished and prevented from increasing do not compensate for the deficit in glucose utilization by up-regulating FFA use.

## INTRODUCTION

Heart failure (HF) is a major and ever-expanding public health challenge, with 1 in 8 deaths in 2019 attributable to HF^1^ despite advances in our understanding of its various etiologies. Whereas etiologies of the syndrome of HF vary widely, a shared characteristic is lack of sufficient myocardial energy supply to sustain continuous contractile activity. Given the obvious and critical importance of this pumping function, energy availability to the heart is finely tuned and tightly regulated: human cardiac muscle consumes 6kg of ATP per day, the majority of which is generated by oxidative phosphorylation in mitochondria^2^. Mature cardiomyocytes in a healthy adult heart derive 60-70% of their energy from free fatty acids (FFA), 20-30% from pyruvate generated from glucose and lactate, and the remainder from branched-chain amino acids and ketones^3^. Studies in mice and humans with HF, however, have demonstrated increased reliance on glucose, mainly from anaerobic ATP production by cytosolic glycolysis rather than from mitochondrial oxidation of pyruvate produced by glycolysis.

The oxygen requirement of FFA oxidation (FAO) per mole of substrate exceeds that of glucose; hence, it has been hypothesized that the diminished supply of oxygen in ischemic HF contributes to the pivot away from FFA as the main energy source^3^. However, this so-called ‘preference’ for glucose for higher cardiac efficiency has also been reported in left ventricular hypertrophy and earlier stages of cardiac dysfunction, conditions in which oxygen availability is thought to be sufficient. In aggregate, multiple discrepancies regarding the roles of various metabolic substrates across different HF etiologies and preclinical models remain unreconciled.

In the case of HFpEF, obesity-driven metabolic syndrome promotes insulin resistance, inhibiting the pivot to glucose utilization as metabolic substrate. Hence, reliance on other substrates is required. Additionally, given the notion that HF is marked by diminished energy supply, important questions arise regarding what substrates are metabolized in HFpEF and whether they fulfill the energy requirements no longer met by glucose.

To address these questions, we have employed a genetic model of cardiac metabolic inflexibility, clamping glucose utilization in the TCA cycle to low levels as seen in HFpEF, to test directly the role of load-induced increases in glucose utilization as energy substrate. We targeted the pyruvate dehydrogenase enzyme complex (PDC), the convergence point and rate-limiting step of glucose and fat metabolism into the Krebs cycle. Mimicking the up-regulation of PDK4 seen in HFpEF^4,5^, we employed a cardiomyocyte-specific transgenic mouse overexpressing the kinase, rendering PDC activity effectively abrogated and blocking glucose oxidation (GO) without manipulating the systemic or local availability of substrates. Using this strategy, we evaluated whether and how metabolic inflexibility affects cardiac function under normal and pressure overload conditions and defined underlying regulatory mechanisms.

## METHODS

The work described here was approved and conducted under the oversight of the UT Southwestern Institutional Animal Care and Use Committee in accordance with the National Institutes of Health (NIH) Guide for the Care and Use of Laboratory Animals.

### Experimental Animals

A PDK4 transgenic (PDK4Tg) mouse line was previously generated to overexpress PDK4 in a cardiomyocyte-specific manner driven by an α-myosin heavy chain promoter^6^. Male founder mice were confirmed to harbor the target sequence by the UT Southwestern Sanger Sequencing Core before being crossed with C57BL/6N female mice (Charles River, Wilmington, MA). Studies described below were conducted in age- and sex-matched mice heterozygous for the PDK4 transgene (PDK4Tg) and their homozygous transgene-negative littermates (WT). Genotype was assessed at 2 weeks of age and confirmed post-mortem via PCR using DNA extracted from tail samples. Animals were housed in a colony room in groups irrespective of their genotype to aid in randomization. The colony room was maintained on a 12:12 light–dark cycle at temperature range of 20-26°C and humidity range of 30%-70%. Animals had *ad libitum* access to water and standard (chow) diet (2016 Teklad Global 16% Protein Rodent Diet, Envigo, Madison, Wisconsin) or a high fat diet (HFD) with 60% kcal from fat (D12492, Research Diets, New Brunswick, NJ).

### Induction of Hypertrophic Remodeling

Male mice were used for studies of cardiac hypertrophic remodeling due to the protective effect of estrogen^7^. Hypertension was induced by addition of L-NAME (L-N^G^-Nitro arginine methyl ester, Sigma-Aldrich) to drinking water at a final concentration of 0.5 g/L at pH=7.4 for six weeks. Blood pressure was measured using the CODA mouse tail-cuff system (Kent Scientific Corporation, Torrington, CT) in awake mice after four days of acclimation to minimize distress. Transverse aortic constriction (TAC) surgery was performed as previously described^8^ on 10–12-week-old mice with minimum body weight of 21g by a surgeon blinded to animal genotype. Negative controls of either genotype underwent sham operations (Sham). As positive controls, WT mice underwent severe TAC (sTAC) surgery^9^, with constriction to 28G instead of 27G, to induce rapid systolic dysfunction. Five to seven days post-surgery, transthoracic echocardiography was performed on conscious, gently restrained mice using a Vevo 2100 system with MS400 transducer. M-mode recordings were performed from a short axis view of the left ventricle at the level of the papillary muscles. Images (triplicate) taken within a heart rate range of 450 to 500 bpm were used for measuring interventricular septal thickness (IVS), left ventricular internal diameter (LVID), and left ventricular posterior wall thickness (LVPW). These measurements were used to calculate left ventricular mass, ejection fraction, and fractional shortening. Echocardiographic studies were performed and analyzed by operators blinded to both genotype and treatment.

### Tissue Collection

At least 24 hours after the last non-invasive evaluation, mice were anesthetized with vaporized isoflurane in a drop chamber and euthanized by cervical dislocation. Unless otherwise noted, tissues used in all experiments were collected between 8am and 9am, ±30min, i.e., ZT2-ZT3, to minimize variations due to circadian rhythm and feeding behavior. Hearts were briefly rinsed in ice-cold Dulbecco’s PBS without calcium or magnesium (Sigma Aldrich) before dissection. Samples were immediately flash frozen by liquid nitrogen submersion and then ground with mortar and pestle for cellular and biochemical assays.

### Molecular Analyses

#### Immunoblot Analysis

Ice-cold RIPA buffer was supplemented with a protease and phosphatase inhibitor cocktail (Roche) immediately before use. Pulverized samples were homogenized in this complete RIPA buffer using a volume of 200-500µL, adjusted to achieve a minimum 2mg/mL protein concentration. Samples were incubated for 30min on ice, sonicated for 1min, and centrifuged for 15min at 13000*g*. The supernatant was collected and filtered through glass wool to eliminate nucleic acid and lipid contaminants. Protein concentration was measured using BCA assay using a commercial kit (Pierce). Samples were diluted in RIPA buffer and Laemmli sample buffer was added. Unless otherwise noted, all samples were denatured by heat for 5min at 95°C. 20µg protein of each processed sample was loaded per well on 4-20% Criterion TGX SDS-PAGE gels (Biorad) for electrophoresis. Proteins in the gel were transferred to 0.45μm nitrocellulose membrane using the Trans-Blot semi-dry rapid transfer system (Biorad). Successful transfer was confirmed by Ponceau red stain. Total protein was measured using Revert-700 stain (Licor) before blocking with fat-free dry milk or BSA in TBS buffer. Incubation times were 12-16h at 4°C for primary antibodies and 1hr at room temperature for secondary antibodies. Antibody list is provided in Appendix 3. Blots were imaged using a Licor Odyssey imaging system and analyzed using Image Studio V5 software.

#### RNA Isolation and qPCR Analysis

RNA was isolated from pulverized tissue using the Total RNA Fatty and Fibrous Tissue Kit (Bio-Rad). A total of 500ng RNA was used for reverse transcription with the iScript cDNA synthesis kit and subsequent real-time qPCR analysis using an CFX instrument and software (Biorad).

### Mitochondrial Assays

#### Mitochondria Isolation

Mice were euthanized by cervical dislocation before rapid excision of hearts perfused with ice-cold isolation buffer containing 10mM MOPS, 210mM mannitol, 70mM sucrose, 1mM EDTA, 20mM NaF, and 1mM DCA. Tissues were homogenized with a POLYTRON disperser (Kinematica, Switzerland) and centrifuged at 650g for 5min at 4°C. Resulting supernatant was filtered through cheesecloth and centrifuged at 10K*g* for 10min at 4°C to obtain pellet containing mitochondria, which was resuspended to 25mg/mL final concentration after measuring protein concentration by BCA method (Pierce, Fisher Scientific).

#### Enzyme Activity Measurements

Isolated mitochondria were diluted to a final concentration of 25µg/mL in room temperature assay buffer containing 25mM MOPS and 0.05% Triton X-100 before loading into a spectrophotometer cuvette (Agilent 8453). Enzyme activity was initiated by addition of 0.1mM CoASH, 0.2mM thiamine pyrophosphate, 1.0mM NAD^+^, 5.0mM MgCl_2_, and the enzyme substrate (1mM pyruvate for PDH, 1mM α-ketoglutarate for α-KGDH, 0.1mM NADH for NADH oxidase). Enzyme activity was calculated from the rate of NAD^+^ to NADH conversion at 340nM.

#### Mitochondrial Respiration Assay

Isolated mitochondria were resuspended in assay buffer containing 10mM MOPS, 210mM mannitol, 70mM sucrose, and 5mM K_2_HPO_4_. For assays using fatty acids as substrate, assay buffer was supplemented with 0.05% BSA. 1mM malate and 0.1mM pyruvate or 0.05mM palmitoylcarnitine were added to the mitochondria before loading into a fluorescence-based oxygen sensor chamber (NeoFox, Ocean Optics). Mitochondrial state 3 respiration was initiated by the addition of 0.25mM ADP after 2min of basal activity at room temperature and was considered completed for purposes of rate calculation when ADP was exhausted.

### Histology

Approximately 2mm thick short-axis sections at mid-papillary muscle level containing both ventricles were dissected at the time of euthanasia and fixed in 4% paraformaldehyde (PFA) for 48 hours at room temperature with agitation. Samples were paraffin embedded and sectioned by UT Southwestern Histopathology Core for H&E, Trichome, picrosirius red, wheat germ agglutinin (WGA), and TUNEL (terminal deoxynucleotidyl transferase dUTP nick end labeling) staining. For each animal, 3 serial sections cut 100 µm apart were imaged using a Zeiss Axiovert100 Inverted Compound Research Photomicroscope with Jenoptik Gryphax Camera for cross-sectional area calculation by WGA. Fifteen total regions of interest, 5 non-overlapping views per section, were captured for each animal to ensure that a minimum of one thousand cardiomyocytes were evaluated (actual cell counts = 2656.2 ± 692). TUNEL-stained sections were imaged in their entirety with software-assisted overlap for automated stitching using a KEYENCE BZ-X700 microscope. A Pacific Image Electronics Prime Histo XE Slide Scanner was used for representative images of non-fluorescent stains. Image processing and quantification of cell cross-sectional area and TUNEL positive nuclei were performed using Image J^10^.

### Targeted Metabolomics Analysis

Left ventricular tissue samples collected 6d ± 0.9 after TAC surgery were homogenized and vortexed in 80% methanol for metabolite extraction. After centrifugation at 20K *g*, the pellet was used to determine sample concentration by BCA assay while the supernatant was lyophilized using a SpeedVac concentrator (Thermo Savant, Holbrook, NY), which was then reconstituted in 0.03% formic acid for analysis by liquid chromatography-tandem mass spectrometry (LC-MS/MS) at the UT Southwestern Metabolomics Facility. A Nexera Ultra High-Performance Liquid Chromatograph system (Shimadzu) and QTRAP 5500 triple-quadrupole mass spectrometer (Applied Biosystems SCIEX, Foster City, CA) were used for metabolite detection as previously described^11^. MultiQuant (AB SCIEX) was used for chromatogram peak area integration which was normalized against the sum of all peak areas detected by the mass spectrometer in that sample (TIC or Total Ion Chromatogram) to obtain relative peak intensity (RPI) and then to the protein content of each sample. Normalized values of 141 detected metabolites for each animal were then evaluated using MetaboAnalyst (https://www.metaboanalyst.ca). Sparse partial least squares discriminant analysis (sPLS-DA) was used to identify metabolites affected by genotype or treatment to maximize separation between the 4 experimental groups. The list of significant metabolites was ranked based on the variable importance in projection (VIP) score. Top metabolites were then mapped to metabolic pathways to calculate their contribution to overall changes in each pathway.

### Protein Pathway Analysis

Mice were euthanized 1 week after TAC, hearts were excised, and left ventricles were dissected. Samples were frozen in liquid nitrogen before being mechanically separated by mortar and pestle to ensure homogeneity. LV samples were then dissolved by sonication on ice in 10mM MOPS buffer with 1mM EDTA at pH=7.4 containing Pierce Protease and Phosphatase inhibitor cocktail (Thermo Fisher Scientific, A32963). Protein concentration was determined by Bradford assay before addition of BSA as a non-endogenous internal standard and 1% SDS. The samples were mixed, heated at 70°C, and precipitated overnight with acetone. The precipitate was reconstituted in Laemmli sample buffer at 1µg/µL and run 1.5cm into an SDS-PAGE gel (short run gel electrophoresis). Each 1.5cm lane was cut as a sample, chopped into smaller pieces, washed, reduced, alkylated, and digested with 1µg trypsin overnight at room temperature. Peptides were extracted from the gel in 50% acetonitrile with 1% acetic acid, and the extracts were taken to dryness by Speedvac and then reconstituted in 200µL 1% acetic acid for analysis.

#### Selected Reaction Monitoring (SRM)

A Thermo Scientific TSQ Quantiva triple quadrupole mass spectrometry system was used in the selected reaction monitoring mode. Specific assay panels were used to measure the respective groups of proteins. Those assays have been tested and validated in prior experiments. The assays monitor two to three peptides per protein. Resolution for both Q1 and Q2 was 0.7 FWHM. A cycle time of 1.5 seconds was used to yield approximately 20 data points across our typical 30sec chromatographic peaks. The LC conditions were a linear gradient elution from 2%B to 45%B in 60min. Total analysis time is approximately 1.5hrs per sample.

#### High Resolution Accurate Mass (HRAM) Analyses

A Thermo Scientific QEXplus instrument was used in the full MS mode, scanning from m/z 300-1100 with a resolution of 280,000 and scan times of 1 scan/second. The LC conditions were a linear gradient elution from 2%B to 45%B in 60min. Total analysis time was approximately 1.5hrs per sample.

#### Data Analysis

All mass spectrometry data were analyzed using the program Skyline. Proper retention times are predicted based on retention time calibration using standard BSA and trypsin peptides. Automated chromatographic peaks assignments by Skyline were edited by manually inspecting the data as needed. The assay for each protein was set to find and integrate 2 to 3 peptides per protein. Calculations determined the total protein response as the geomean of the monitored peptides. Results are normalized to the BSA internal standard and expressed as pmol/100µg total protein.

### ^13^C Nuclear Magnetic Resonance Isotopomer Analysis of Substrate Metabolism

Five days after TAC, mouse hearts were excised, cannulated via the aorta, and connected to a perfusion column apparatus maintained at 37⁰C using a controlled temperature bath. Hearts were retrograde perfused for 30min at 100cm-H_2_O pressure with a modified Krebs-Henseleit buffer containing 8 mM [1,6-^13^C]glucose, 1.2 mM [2-^13^C]lactate, 0.12 mM [2-^13^C]pyruvate, 0.4 mM [U-^13^C]long chain fatty acids (LCFA) with 0.75% bovine serum albumin (BSA) and 50µU/mL insulin. The non-recirculating buffer was oxygenated with a thin-film oxygenator with a 95:5 mixture of O_2_:CO_2_. Heart rate was monitored throughout the perfusion with a fluid-filled catheter in the left ventricle. Oxygen consumption was measured by collecting coronary flow samples into a gas-tight syringe at 5 and 25 min and analyzed using a blood gas analyzer (Instrumentation Laboratory, Lexington, MA). Hearts were snap-frozen immediately after 30 min of perfusion, pulverized in liquid nitrogen and extracted with perchloric acid (4%). The perchloric acid extract was then neutralized and reconstituted in D_2_O containing 0.5 mM 2,2-dimethyl-2-silapentane-5-sulfonate (DSS) standard. Proton-decoupled ^13^C-NMR spectra of heart extracts were acquired using a 14.1T spectrometer (Bruker Corporation, USA) equipped with 5-mm cryoprobe. ^13^C-NMR multiplets from glutamate were deconvoluted using ACD/SpecManager (ACD Labs, Canada) and multiplet ratios were used to determine the relative oxidation of [1,6-^13^C_2_]glucose, [2-^13^C]lactate-[2-^13^C]pyruvate, [U-^13^C]FA, and unlabeled endogenous substrates (e.g., triglycerides and glycogen). Fractional areas of ^13^C multiplets from the fitting were used for estimating PDC flux relative to that of citrate synthase (CS) using flux analysis software tcaCALC^12^. Absolute fluxes of CS and PDH were calculated according to a previously reported method^13^. Glycolysis was measured by ^1^H-NMR of effluent to detect conversion of [1,6-^13^C] glucose to [3-^13^C] lactate.

### Statistical Analysis

Statistical analyses were performed using GraphPad Prism v.9 (GraphPad Software, Inc, San Diego, USA) unless otherwise indicated above. All data are presented as mean ± SEM. Statistical significances were tested using 2-way analysis of variance (ANOVA) with Tukey correction for *post hoc* multiple comparisons. Differences were considered significant at ≤ 0.05 and indicated with asterisks on plots corresponding to number of digits after the decimal.

## RESULTS

To elucidate the role of metabolic flexibility in HFpEF pathophysiology, we evaluated the myocardial response to disease-related stress by blocking pyruvate utilization within the Krebs cycle with a transgenic mouse model of PDK4 overexpression (PDK4Tg) driven by the αMHC gene promoter^6^. By design, the PDK4 transgene was transcribed abundantly **(<u>Fig S1A, S1B</u>)** without altering expression of respiratory chain complexes **(<u>Fig S1C</u>)**, suggesting specificity. As expected, this led to phosphorylation of PDC specifically at site 1 (human Ser264/mouse Ser293) **(<u>Fig S1D</u>)**. Consistent with the fact that phosphorylation of just 1 of the 6 available serine residues in its E1α subunit is sufficient to inactivate PDC^14^, we observed a 20-fold decline in PDH enzymatic activity in mitochondria isolated from PDK4Tg hearts **(<u>Fig S1E</u>)**. This decline in PDH activity was independent of oxidative stress-induced enzyme instability, as addition of DTT *in vitro* had no effect (**<u>Fig S1E</u>)**. Activity of alpha-ketoglutarate dehydrogenase (α-KGDH), another Krebs cycle enzyme harboring the E1α subunit, was not affected by PDK4 overexpression, again pointing to specificity of PDK4 interaction with PDC **(<u>Fig S1F</u>)**. We also found that ADP-dependent respiratory rate in mitochondria isolated from PDK4Tg mice was similar to WT mice when supplied with palmitoyl carnitine (PC) as respiratory substrate, yielding a respiratory control ratio (RCR) of 8 in WT mice vs 8.3 in PDK4Tg (**<u>Fig S1G</u>**). Due to lower state 3 respiratory activity in PDK4Tg mouse mitochondria, RCR using pyruvate was 3.9 in WT vs 1.9 in PDK4Tg, indicating mitochondrial respiratory dysfunction and inefficiency in the latter (**<u>Fig S1G</u>**). These findings confirmed that cardiomyocyte-specific PDK4 overexpression is sufficient to essentially eliminate PDC activity without altering other metabolic processes, lack of which precludes use of pyruvate for ATP generation through the Krebs cycle.

HFpEF myocardium is marked by metabolic inflexibility^15^. Having documented suppression of metabolic flexibility in our PDK4Tg hearts, we then evaluated their ability to accommodate enhanced energy demand elicited by modest increases in afterload. As prior work tested the effects of an overwhelming trigger of cardiac remodeling, calcineurin transgenesis^6^, we employed a milder, more physiologically relevant, degree of stress triggered by L-NAME. Adult male PDK4Tg mice and their WT littermates (9.3wk ± 1) were exposed to L-NAME, an inhibitor of constitutive NO synthases, which has been shown previously to induce hypertension and mild diastolic but not systolic dysfunction with long term exposure^16^. As expected, animals treated for 6 weeks with 0.5g/L L-NAME manifested increased systolic and diastolic blood pressure (BP) (**Fig 1A**). Echocardiography revealed no changes in systolic function of the WT hearts exposed to L-NAME treatment **(Fig 1B, <u>Table S1</u>)**. In contrast, PDK4Tg mice exposed to L-NAME manifested significantly lower LV % fractional shortening (**Fig 1B**) and higher end-systolic LV volumes (**Fig 1C**), consistent with diminished myocardial contractility. Hearts of PDK4Tg mice exposed to L-NAME manifested a dilated phenotype with thinning of the left ventricular wall (**Fig 1D**). In contrast, contractile dysfunction emerged at a point prior to significant hypertrophic growth; whereas LV mass by echo trended higher in both groups on exposure to L-NAME, it did not achieve statistical significance (**<u>Table S1</u>)**. In addition, heart weights of L-NAME-treated mice were not greater than controls (**Fig 1E)**. Based on these findings, we concluded that PDK4Tg mice manifested early declines in contractile function in response to modestly increased afterload without accelerated hypertrophic remodeling.

**Figure 1.**
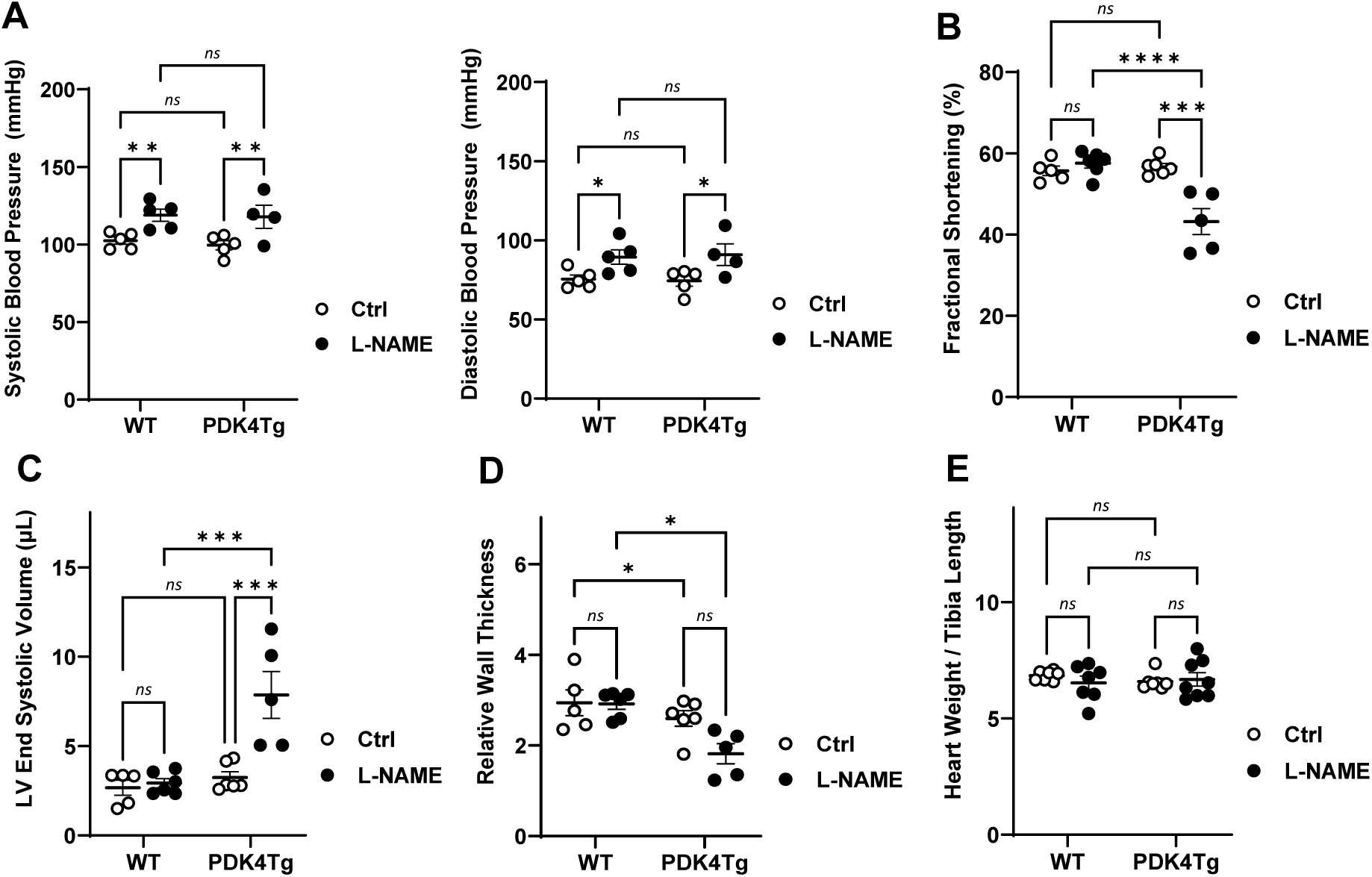
Metabolic flexibility is required to maintain cardiac contractile function in hypertension. A. Tail vein blood pressure (SBP, DBP) measurements of PDK4Tg mice and their WT littermates after consuming water containing 0.5g/L L-N^G^-nitro arginine methyl ester (L-NAME) or not (Ctrl) for 6 weeks. Quantitative analyses for echocardiographic evaluation of left ventricular (LV) % fractional shortening (FS), B., LV end systolic volume (LVESV), C., and LV relative wall thickness, D. E. Quantitative analysis of post-mortem morphometric evaluation of heart size after 7 weeks of L-NAME treatment. Statistical significance was determined by 2-way ANOVA with Tukey’s post hoc MCT. Asterisks indicate number of zeroes in p values of post-hoc multiple comparisons: *= p≤.05, **= p≤.005, ***= p≤.0005, ****= p≤.00005.

L-NAME-induced hypertension is mild and develops gradually over several weeks. In light of this, we subjected PDK4Tg mice and their WT littermates to transverse aortic constriction (TAC) surgery to trigger a local and acute increase in afterload. As early as 5 days after TAC surgery, PDK4Tg mice manifested robust decreases in LV % fractional shortening as compared with sham-operated animals, changes not seen with WT littermates (**Figs 2A, 2B**). Similar to L-NAME treatment, TAC provoked dilation of the LV chamber (**Figs 2A, 2C**) and thinning of the LV wall (**Fig 2D**), which were substantially more severe in PDK4Tg hearts (**<u>Table S2</u>**). This acute failure phenotype mimicked that observed with the less robust afterload intervention, L-NAME treatment, confirming the contractile dysfunction is due to the modestly increased afterload and not systemic effects.

**Figure 2.**
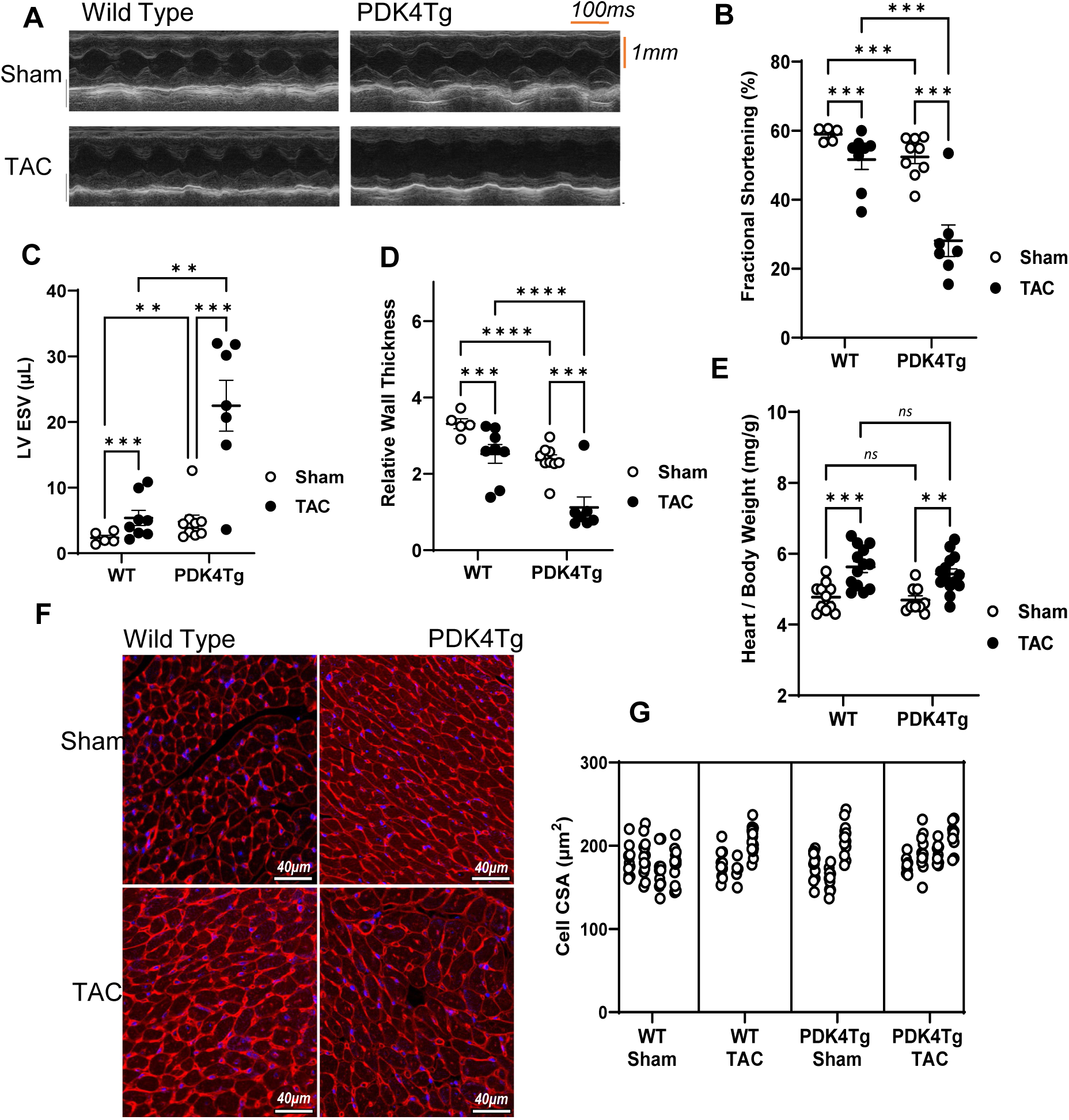
Metabolic flexibility is required to maintain cardiac contractile function in response to acute increase in afterload. A. Representative M-mode echo images of WT and PDK4Tg mice 6 days after surgery. Quantitative analyses for echocardiographic evaluation of left ventricular (LV) fractional shortening (FS), B., LV end systolic volume (LVESV), C., and LV relative wall thickness, D. E. Quantitative analysis of post-mortem morphometric evaluation of heart size 8 days after surgery. F. Representative images and G. quantification of cardiac myocyte cross-sectional area by wheat germ agglutinin (WGA) staining. Scale bar = 40µm. Technical replicates shown by animal per column; each count represents mean cell area per view of at least 100 cells for that animal. Statistical significance was determined by 2-way ANOVA with Tukey post hoc MCT. Asterisks indicate number of zeroes in p values of post-hoc multiple comparisons: *= p≤.05, **= p≤.005, ***= p≤.0005, ****= p≤.00005.

### Hypertrophic remodeling does not drive contractile dysfunction in PDK4Tg hearts

Our initial results suggested that acute imposition of moderately elevated afterload with resultant myocardial contractile dysfunction in PDK4Tg transgenic mice occurs independently of an exacerbated hypertrophic response. To test this further, we compared myocardial hypertrophy and hypertrophic signaling pathways in WT and PDK4Tg hearts. The extent of myocardial hypertrophic growth in WT mice 5 days after TAC surgery evaluated by echocardiography (**<u>Table</u> <u>S2</u>**), at 1-week post-TAC evaluated by morphometric analysis (**Fig 2E**), and on exposure to substantially greater afterload stress (sTAC)^9^ (**<u>Fig S2A</u>**) were each greater than the same measures of hypertrophic remodeling with L-NAME in each genotype. WGA (wheat germ agglutinin) staining of LV tissue samples collected at mid-papillary muscle level were grossly similar **(Fig 2F)**, and quantification of cell cross-sectional area revealed no difference in cellular hypertrophy due to TAC treatment or PDK4 overexpression (**Fig 2G**).

Immunoblot analysis of proteins involved in hypertrophic pathways driven by mTOR and MAPK in the LV revealed similar findings. Phosphorylation of p70-S6 kinase was increased to a similar degree after TAC in both groups (**Fig 3A)**, whereas ERK1/2 was upregulated to a greater extent in PDK4Tg hearts (**Fig 3B**). However, this was not sufficient to activate NFAT, as RCAN1.4 abundance increased after TAC to the same extent in both groups (**Fig 3C**). In addition, JNK1/2 phosphorylation levels were unchanged, suggesting that competitive JNK inhibition of NFAT did not mask the effect of ERK1/2 (**Fig 3D**).

**Figure 3.**
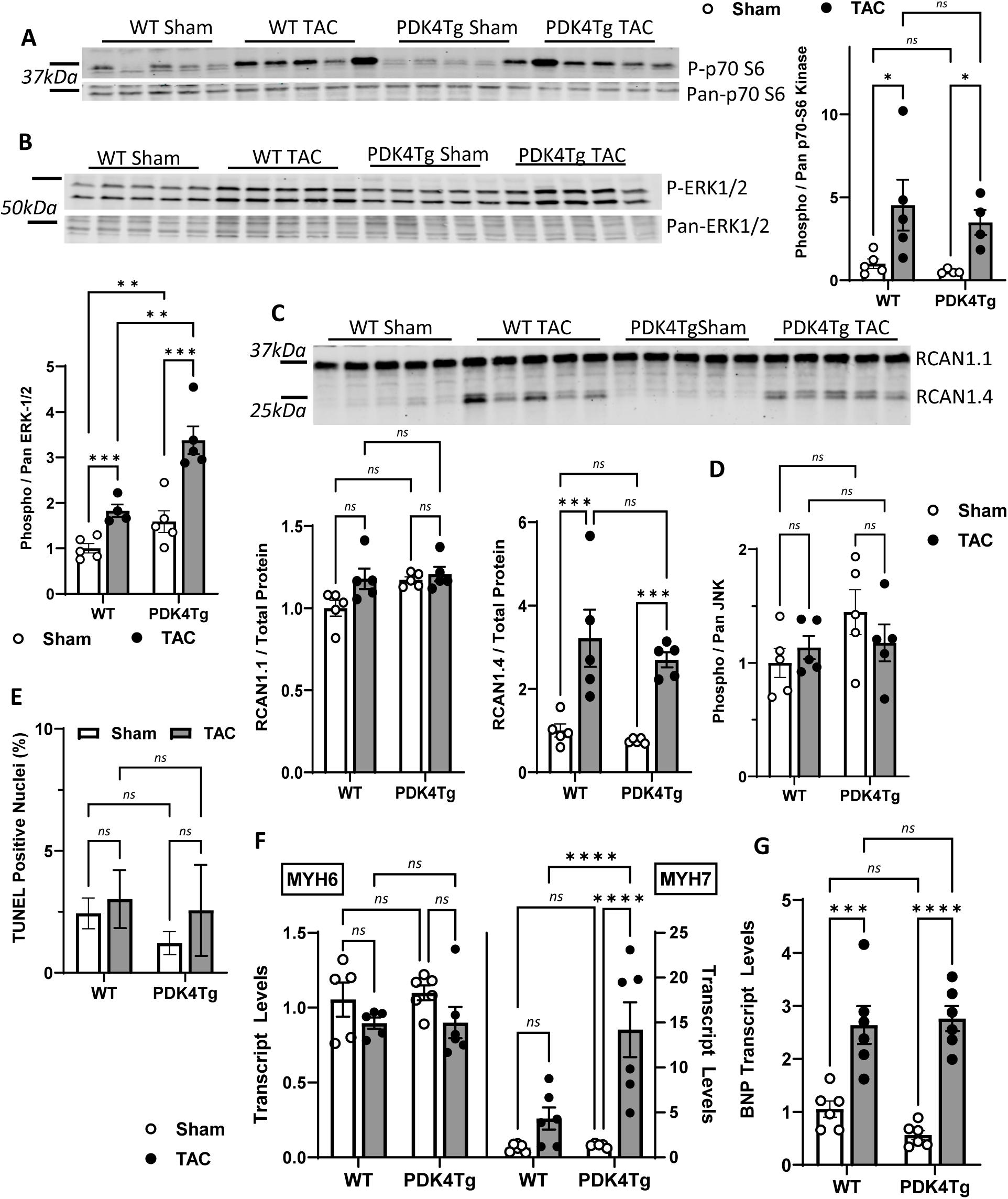
Afterload-induced hypertrophic signaling does not drive contractile dysfunction in the setting of metabolic inflexibility. A-D. Representative immunoblot and quantification of proteins in hypertrophic signaling pathways downstream of mechanical stress 1 wk after surgery in LV samples from WT and PDK4Tg mice (n=5). A. p70-S6K & B. ERK1/2 activation determined by ratio of phosphorylated to total protein. C. Regulator of calcineurin 1 (RCAN1) isoforms 1 (RCAN1.1) and 4 (RCAN1.4) normalized to total protein abundance per lane (data not shown) D. JNK1/2 activation assessed by ratio of phosphorylated to total protein (n=5). Statistical analysis was performed on fold change of each group over WT sham control. E. TUNEL (terminal deoxynucleotidyl transferase dUTP nick end labeling) staining in WT and PDK4Tg mice 1wk after surgery (n=3-4). Real-time quantitative PCR (RT-qPCR) measurement of F. alpha- and beta-myosin heavy chain and G. B-type natriuretic peptide (BNP) mRNA levels in WT and PDK4Tg mice LV tissue 1wk after surgery (n=5-6). Asterisks indicate number of zeroes in p values of Tukey post-hoc multiple comparison tests (MCT) in 2-way ANOVA: *= p≤.05, **= p≤.005, ***= p≤.0005, ****= p≤.00005.

Earlier work suggested that PDK4 over-expression could trigger cell death^6^. To test for this, we evaluated the possibility that apoptotic cell death eliminated rapidly hypertrophying cells from the sampled tissues: TUNEL staining (**Fig 3E**) and caspase 3 cleavage (**<u>Fig S2B</u>**) levels revealed no differences between WT and PDK4Tg mouse LVs. H&E staining of LV sections from these mice also revealed no signs of necrosis or fibrosis (**<u>Fig S2C</u>**).

We next investigated the effects of hypertrophic signaling on gene transcription. Interestingly, whereas we detected no significant decrease in αMHC levels in either group, levels of βMHC were significantly greater in PDK4Tg hearts after TAC when compared with WT TAC controls (**Fig 3F**). Since this increase in βMHC mRNA in the setting of PDK4 overexpression was not observed in sham-treated animals, it arises as a component of the response to afterload stress. In line with our Western blot findings, BNP transcript levels were increased to comparable degrees in the ventricles of both genotypes (**Fig 3G**). Collectively, these data led us to conclude that the hypertrophic signaling cascade elicited in response to increased afterload, as well as the subsequent remodeling process, were not altered by cardiomyocyte overexpression of PDK4.

### Oxidative stress due to increased FFA utilization does not account for contractile dysfunction in PDK4Tg hearts

These findings, coupled with previous work^6,17^, revealed limited cardiomyocyte utilization of pyruvate in αMHC-PDK4Tg mice with presumed increases in fractional oxidation of long-chain fatty acids (LCFAs). As oxidation of fatty acids produces more reactive oxygen species (ROS) than glucose^18^, we theorized that increased reliance on fatty acids might elicit oxidative stress in cardiomyocytes. NADH oxidase enzyme activity measured in mitochondrial isolates from PDK4Tg mice manifested no significant differences (**Fig 4A)**, indicating lack of uncoupling that would trigger higher ROS production in the electron transport chain. We also observed lower levels of protein carbonylation in LV lysates of PDK4Tg mice one week after sham or TAC surgery, yet TAC did not elicit a statistically significant change in either genotype (**Fig 4B**). These findings provided the first hint that increased FAO was not occurring in the PDK4Tg mice and suggested that they have developed compensatory mechanisms for handling persistently increased oxidative stress.

**Figure 4.**
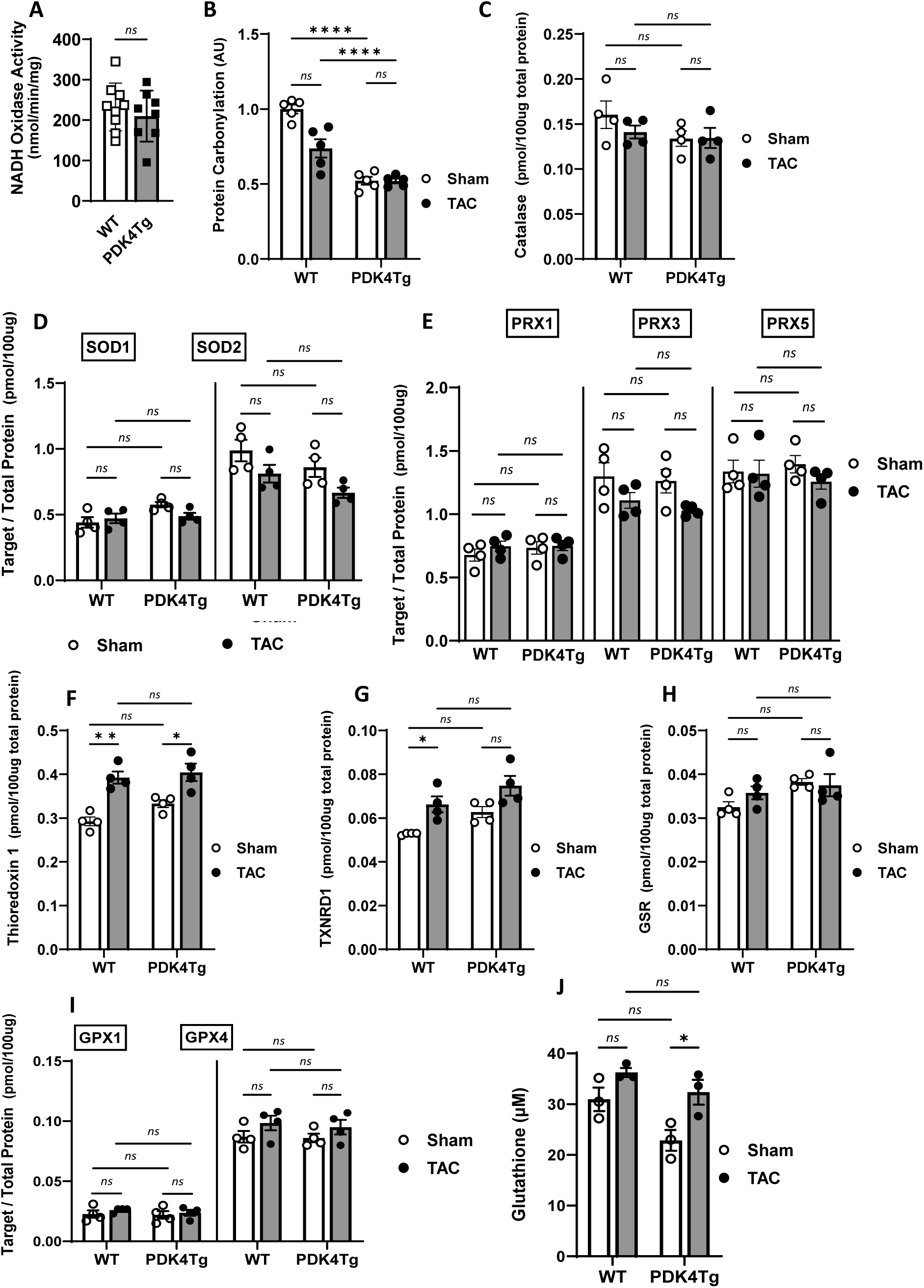
ROS increases due to PDC inhibition are insufficient to cause contractile dysfunction. A. NADH oxidase activity measured by spectrometry in mitochondria isolated from hearts of WT and PDK4Tg mice at 10-12 weeks of age. Statistical analysis by unpaired t-test, ns = p>.05. B. Calorimetric quantification of protein carbonylation in PDK4Tg and WT LV tissue 7d after surgery by tagging DNP hydrazones. C-I. Absolute amounts of ROS handling proteins as labeled in graph titles, as measured by selective reactive monitoring (SRM) mass spectrometry. All performed simultaneously on the same LV samples collected from PDK4Tg and WT mice 7d after surgery, n=4. SOD: superoxide dismutase, TXNRD: thioredoxin reductase, GSR: glutathione-disulfide reductase, GPX: glutathione peroxidase. J. Calorimetric quantification of total glutathione in WT and PDK4Tg LV samples 7d post-surgery via its reaction with GSR and Ellman’s Reagent. Statistical analysis was performed by 2-way ANOVA for genotype and surgery main effects model with Tukey post-hoc multiple comparisons for B-J. Significance threshold set to .05 with following labels *= p≤.05, **= p≤.005, ***= p≤.0005, ****= p≤.00005.

To corroborate this, we measured absolute levels of several proteins known to handle ROS in the heart by means of selective reaction monitoring (SRM). We observed no difference in the levels of cytosolic catalase (**Fig 4C**) or superoxide dismutase 1 (SOD1) across all groups (**Fig 4D**), suggesting unchanged cytosolic handling of hydrogen peroxide (H_2_O_2_) shuttled out of mitochondria. Similarly, we observed no differences in the levels of the mitochondrial proteins superoxide dismutase 2 (SOD2) (**Fig 4D**) or peroxiredoxins (PRX) 1, 3, and 5 (**Fig 4E**), whereas both SOD2 and PRX3 were slightly but significantly decreased by TAC treatment (2-way ANOVA, p=0.0178, p=0.0203, n=4). As these proteins neutralize H_2_O_2_ within mitochondria, these results taken together with our findings with cytosolic ROS scavengers indicate that H_2_O_2_ production is unlikely to be affected by PDK4 overexpression.

NADPH is a substrate critical to energy homeostasis, decreased abundance of which induces pathological hypertrophy and apoptosis^19^. Proteins involved in maintaining oxidative/reductive balance also govern PRXs and glutathione. We again employed SRM on LV from 12-week-old PDK4Tg mice and their WT littermates to investigate the effect on these proteins of TAC-induced increases in afterload. We observed TAC-induced (1 week) increases in both thioredoxin (TXN, p<0.0001) and TXN-reductase (TXNR, p=0.0016 by 2-way ANOVA, n=4) in both genotypes (**Fig 4F<u>, 4G</u>**). In contrast, levels of glutathione-disulfide reductase (GSR) which catalyzes the complementary reaction to TXNR1 were unaffected by TAC (**Fig 4H)**, p=0.4652 by 2-way ANOVA, n=4). We also observed no difference in the levels of glutathione peroxidase (GPX) 1 or 4 (**Fig 4I**).

The lack of change in the levels of GPX1, GPX4, and GSR suggests lack of alteration in the reductive balance of glutathione, which would indicate changes in TXN and TXNR levels related to their role in NADPH homeostasis alone. Although we were unable to directly evaluate the levels of oxidized and reduced glutathione for comparison, total glutathione levels followed the pattern observed above: a robust increase elicited by TAC (p=0.0063) in both WT and PDK4Tg mice (**Fig 4J**). With respect to total glutathione abundance, 2-way ANOVA revealed a decrease in the abundance in PDK4Tg mice, as well (p=0.0176). No interaction was observed between genotype and treatment, suggesting that PDK4Tg mice were able to produce more glutathione, similar to WT littermates, in response to TAC. This suggests that persistent ROS in PDK4Tg hearts is insufficient to exhaust their capacity for glutathione production. Collectively, these findings indicate that the slight increase in ROS due to PDK4 overexpression is surpassed by the effect of TAC and is not solely responsible for the severe systolic contractile dysfunction observed in PDK4Tg animals after only 5 days of TAC.

### Fractional oxidation of FFA decreases after TAC, independent of altered glucose use

As described above, ROS levels are not robustly increased in PDK4Tg hearts exposed to increased afterload. This is surprising given that expected increases in fat oxidation in these hearts would likely elicit increased ROS generation. As we had confirmed complete block of pyruvate use in cardiomyocytes by PDC inhibition, these findings raised the possibility of enhanced contribution to energy production from other, normally scarcely utilized, alternative substrates such as glycogen, amino acids, and ketones. To test this, we used NMR isotopomer analysis *ex vivo* in the hearts of PDK4Tg mice and their WT littermates 5 days after TAC. Physiological levels of ^13^C-labeled glucose, pyruvate, lactate, and long-chain fatty acids (LCFA) were delivered to unloaded hanging hearts. Consistent with previous reports^20^ ^21^, pressure overload-driven metabolic remodeling led to diminished utilization of lipids for oxidative energy production in both WT and PDK4Tg animals (**Fig 5A**). Whereas WT hearts were able to upregulate glucose, lactate, and pyruvate use to compensate for this change, PDK4Tg hearts were not, but rather relied more heavily on unlabeled endogenous substrates such as glycogen, amino acids, or ketones. As expected, both WT and PDK4Tg mice increased oxygen uptake after TAC given the requisite enhanced mitochondrial oxidative respiration rate (**Fig 5B**). Surprisingly, PDK4Tg mice manifested 30% lower oxygen demand at baseline than their WT littermates, and this difference was preserved after TAC (2-way ANOVA followed by Tukey’s multiple comparison test (MCT), p<0.0001). Consequently, when the absolute oxidation rate of each substrate was calculated from the above, we found that the amount of LCFA substrate used for energy production was not higher in PDK4Tg animals as we had expected (**Fig 5C**). Coupled with the small decrease in LCFA utilization after TAC in both groups (2-way ANOVA, p=0.0175), PDK4Tg mice did not manifest increased capacity for fat utilization to maintain TCA cycle flux (**Fig 5D**). Whereas PDK4Tg mice were able to increase TCA flux after TAC to a degree similar to that of WT mice (WT=12.84% vs PDK4Tg=12.44%), these levels were substantially lower than those in WT hearts after either sham or TAC surgery (2-way ANOVA, p<0.0001 for both genotype and surgeries). This was due to robust inhibition of PDH flux elicited by over-expression of PDK4 (p<0.0001), which prevented PDC activation after TAC (mean:0.5638 vs 0.9105, not statistically significant) in contrast to WT mice (sham = 1.972 vs TAC = 4.075, p<0.0001, 2-way ANOVA with Tukey’s post hoc MCT in each instance) (**Fig 5E**).

**Figure 5.**
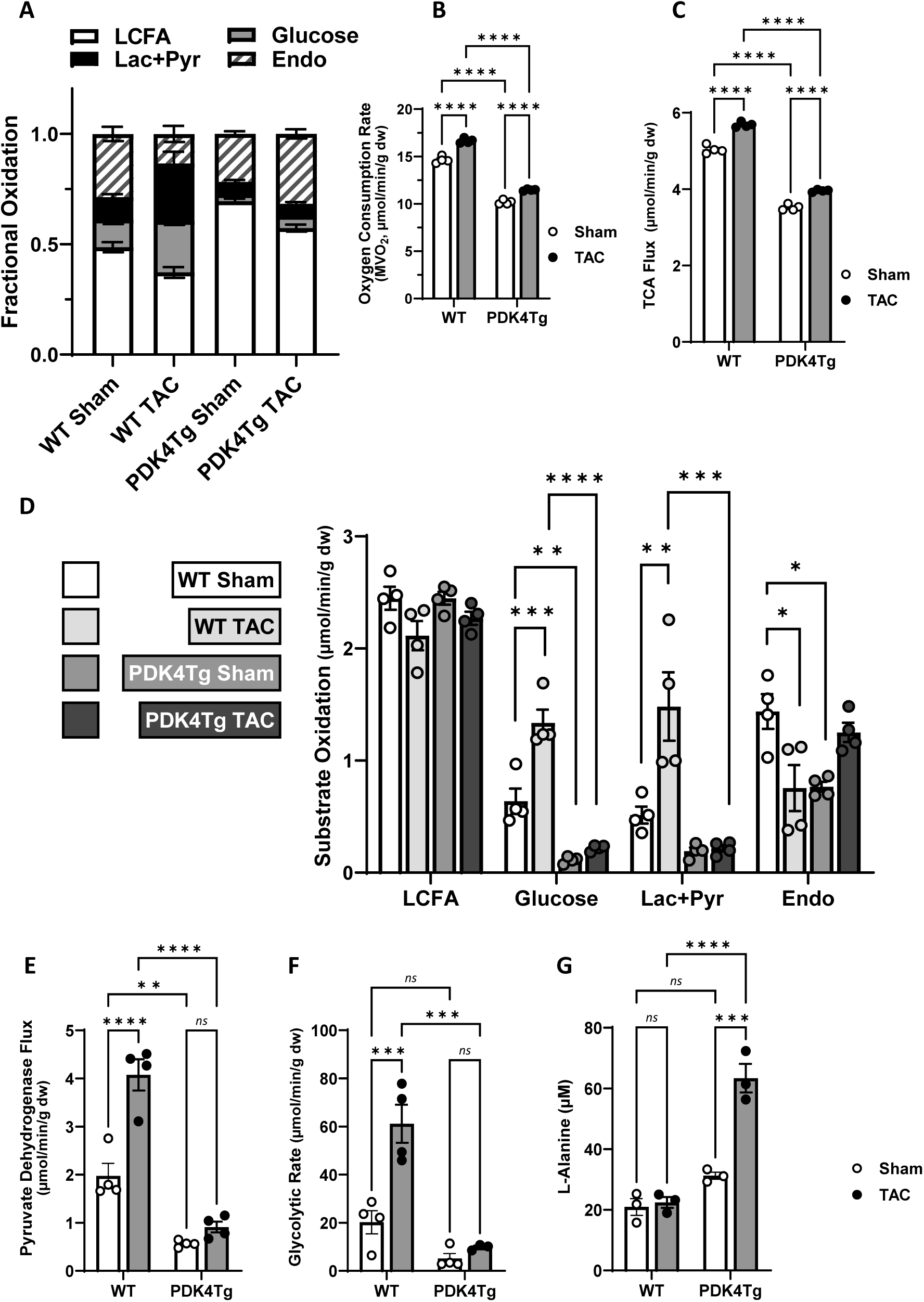
FAO is not adequately upregulated after PDC inhibition to maintain contractile function. A. Relative oxidation of different substrates in WT and PDK4Tg hearts ex vivo by NMR isotopomer analysis 5 days post-TAC (n=4). LCFA: long chain fatty acids, Lac + Pyr: lactate and pyruvate, Endo: endogenous unlabeled substrates. B. Oxygen consumption rate by blood gas analysis in WT and PDK4Tg hearts ex vivo 5 days post-TAC, normalized to final total heart weight. C. Absolute levels of oxidation of the ^13^C labeled exogenous substrates and endogenous unlabeled substrates in WT and PDK4Tg hearts ex vivo 5 days post-TAC, grouped by substrate. D. TCA cycle flux and E. PDH enzyme flux calculated using tcaCALC software to find best fitting models of relative substrate oxidation. F. Glycolytic rate in ex vivo hearts determined by the conversion of labeled glucose to lactate over 30 min, normalized to dry heart weight. G. Concentration of L-alanine in LV samples of WT and PDK4Tg mice 7d post-TAC, measured by the color intensity of alanine transaminase enzyme product at 570nm. Statistical significance of genotype and surgery determined by 2-way ANOVA for each substrate with a main effects model with Tukey post-hoc comparisons between individual groups. Only significant group comparisons of p≤.05 are listed. *= p≤.05, **= p≤.005, ***= p≤.0005, ****= p≤.00005.

In the setting of hypoxia, such as *in utero* or during myocardial ischemia, cardiomyocytes rely on cytosolic glycolysis relative to mitochondrial oxidative phosphorylation for energy production^22^. As pyruvate and lactate, products of glycolysis, are in cellular equilibrium, we theorized that the abundance of unused pyruvate, and thus lactate, in PDK4Tg hearts would inhibit glycolysis. Indeed, measurements of labeled lactate in the myocardial effluent, produced by glycolysis from exogenous glucose, supported this hypothesis as genotype accounted for 46.9% of the variation in glycolic rate (2-way ANOVA, p<0.0001) (**Fig 5F**). Pressure overload, on the other hand, triggered upregulation of glycolysis in both WT and PDK4Tg hearts (p=0.0011, 22.13% of total variation), albeit to a lesser extent in the PDK4Tg mice (PDK4Tg sham vs TAC not statistically significant, 2-way ANOVA with Tukey’s post hoc MCT in each instance) (**Fig 5F**).

Based on these findings, we concluded that PDH inhibition by PDK4 not only blocks upregulated glucose utilization by the TCA cycle, it also blunts glycolysis in the cytoplasm. As pyruvate can be converted to L-alanine^23^, we measured the levels of this amino acid in the myocardial effluent. Consistent with the notion that excess pyruvate accumulating in the cytosol is converted to alanine, we detected a 50% increase in L-alanine concentration measured in PDK4Tg LV samples 1 week after sham and over 200% after TAC (2-way ANOVA, 52.84% variation due to genotype, p<0.0001) (**Fig 5G**).

### Decreased FAO enzyme abundances independent of increased glucose/pyruvate utilization

Given the surprising lack of increased fatty acid utilization in PDK4Tg mice exposed to TAC, we next evaluated FAO and glucose utilization pathways. RT-qPCR analysis of ventricular lysates of PDK4Tg mice and their WT littermates 1 week after TAC revealed diminished abundances of transcripts coding for acyl-CoA dehydrogenase enzymes (ACAD) for short-, medium-, and very long-chain FFAs compared to sham animals irrespective of genotype (2-way ANOVA, ACADs p=0.0071, ACADm p<0.0001, ACADvl p=0.0101) (**Fig 6A**). We also subjected WT mice to a more severe form of TAC (sTAC)^9^ to induce rapid systolic dysfunction to a degree similar to that observed with PDK4Tg mice after TAC. This sTAC group with HF manifested a more pronounced decrease in the abundance of ACAD transcripts, which was not statistically different from either WT TAC or PDK4Tg TAC groups. Based on this, we conclude that these changes are not a consequence of HF per se but rather elements within the progression of systolic dysfunction.

**Figure 6.**
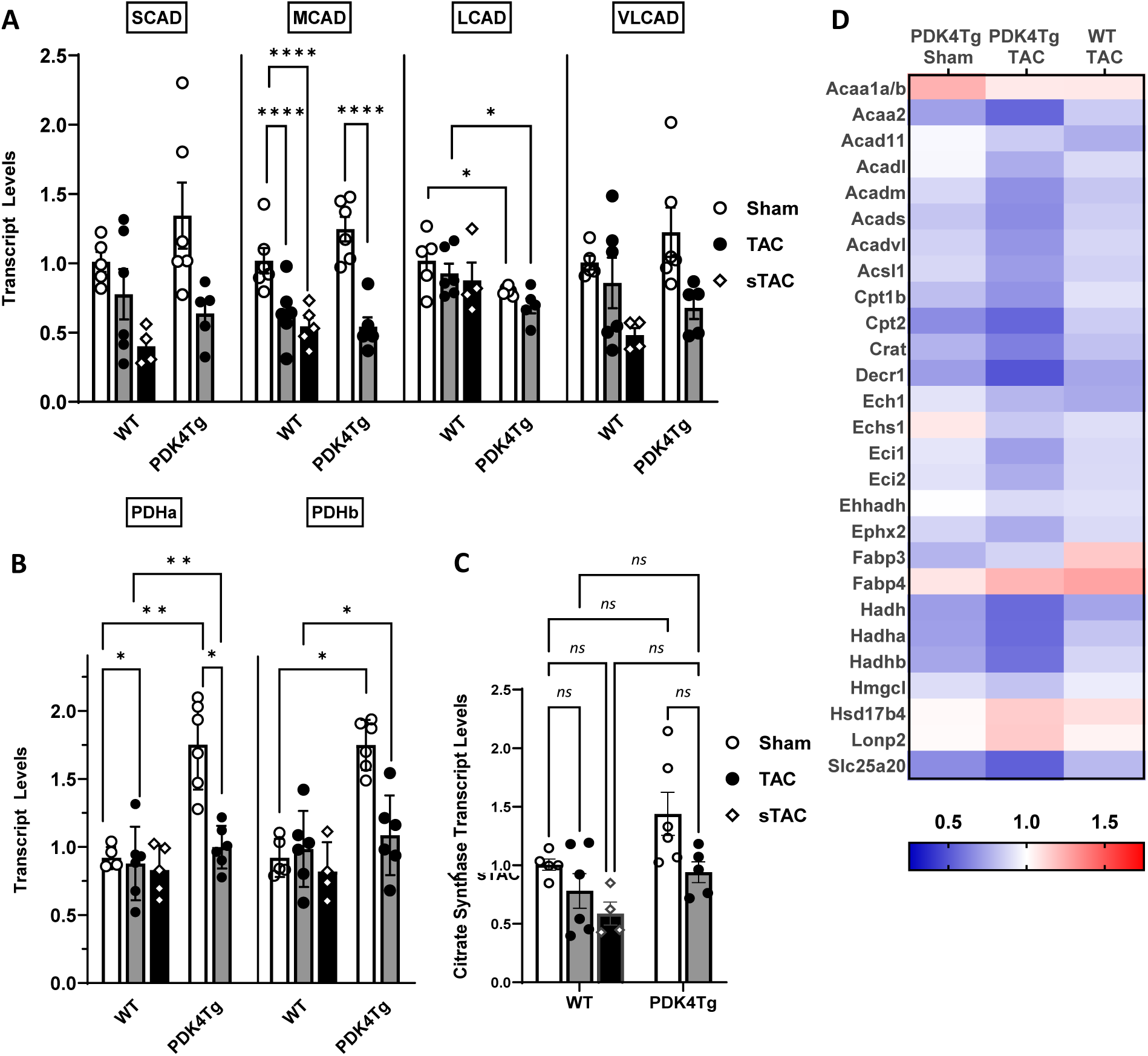
FAO pathways are paradoxically downregulated by PDK4 overexpression. Expression of enzymes and transporters involved in glycolysis, fatty acid beta oxidation, and TCA cycle were measured at transcript and protein level in LV samples collected from PDK4Tg mice and their WT littermates 7 days post-surgery. All results are presented as fold change of each group over WT sham controls. A. mRNA transcript levels of acyl-CoA dehydrogenase enzymes (ACAD). SCAD: short-chain FA ACAD, MCAD: medium-chain FA ACAD, LCAD: long-chain FA ACAD, VLCAD: very-long-chain FA ACAD. TAC = transverse aortic constriction, sTAC = severe TAC. Comparisons shown for only statistically significant results as determined by Tukey MCT after 2-way ANOVA. B. mRNA transcript levels of pyruvate dehydrogenase (PDH) E1 component subunits alpha (PDHa) and beta (PDHb). Comparisons shown for only statistically significant results as determined by Tukey MCT after 2-way ANOVA. C. mRNA transcript level of citrate synthase enzyme. For all above, asterisks indicate number of zeroes in p values of Tukey post-hoc multiple comparison tests (MCT) in 2-way ANOVA: *= p≤.05, **= p≤.005, ***= p≤.0005, ****= p≤.00005. *ns* = not significant, p>.05. D. Visualization of beta oxidation pathway analysis for proteins found to be statistically significantly changed by full model 2-way ANOVA. Each column represents an experimental group’s mean value for a protein per row. Colors represent the amount of change over WT sham for the other 3 groups. WT sham mean value for each protein set to 1. Blue: <1, less than WT sham. Red: >1, more than WT sham. Range of color scale determined by max and min values of data as .25 – 1.75. Full protein names can be found in Table S3.

Several other transcripts involved in the transport and oxidation of FFA were also downregulated after TAC in both groups (**Fig 6A, <u>Table S3</u>**). Of note, mRNA levels of long-chain acyl-CoA dehydrogenase (ACADl) differed by genotype, with PDK4Tg mice manifesting slightly lower expression than their WT littermates (p=0.0036). We also found that genes encoding PDH E1 subunits (**Fig 6B**) and the TCA cycle enzyme citrate synthase (**Fig 6C**) were downregulated after TAC but upregulated in PDK4Tg mice, culminating in comparable mRNA levels in WT sham and PDK4Tg TAC hearts.

We then confirmed that the downregulation of FFA oxidation transcripts triggered by TAC led to lower levels of the corresponding proteins. SRM replicated the trends observed with the transcripts probed above as additional proteins involved in the transport and processing of FFAs in mitochondria, albeit to a lesser extent than that seen with mRNA levels (**Fig 6D**). Furthermore, we detected a greater impact of PDK4 on the downregulation of the majority of these proteins, which was not statistically significant at the mRNA level, likely due to the higher sensitivity of the protein detection methods. The difference between protein and mRNA regulation may also be due to an additional control mechanism, such as decreased mRNA stability or increased protein degradation. On the other hand, proteins in the glycolysis pathway (**<u>Fig S3A</u>**), TCA cycle (**<u>Fig</u> <u>S3B</u>**), and electron transport chain (**<u>Fig S3C</u>**) were significantly changed (2-way ANOVA) and were more often altered in opposite directions by PDK4 overexpression versus TAC treatment, consistent with our NMR findings.

## DISCUSSION

The myocardium is capable of pivoting across multiple metabolic substrates to generate the ATP required for continuous mechanical contraction. In the setting of disease-related stress, the heart pivots toward increased metabolism of glucose; this response, however, is greatly attenuated in the settings of high fat diet exposure or diabetes as is typical of cardiometabolic HFpEF. Here, we set out to define the role of this HFpEF-attenuated swing toward glucose utilization absent the exogenous effects of lipid over-exposure by clamping glucose utilization to near zero by suppressing PDC via transgenic over-expression of PDK4. We report that the ventricular response to modest elevations in afterload in this circumstance is marked by rapid contractile dysfunction. Hypertrophic growth, however, is unaltered. Under resting conditions, the myocardium, as expected, relied on other, non-glucose substrates. Surprisingly, however, in the setting of afterload stress, the myocardium, depleted of its access to pyruvate as a metabolic substrate, does not increase utilization of FFA; rather, FAO is paradoxically diminished, and utilization of other substrates, such as amino acids and ketone bodies, increases.

### Myocardial metabolism in cardiovascular diseases

Abnormalities of energy metabolism have been described in a wide range of clinical conditions that affect cardiovascular health, including but not limited to hypertension, obesity and diabetes, heart failure, and cardiomyopathies due to inborn errors of metabolism^22^. Therefore, elucidating the contribution of cardiac metabolic derangements to the pathogenesis of heart disease, including HFpEF, remains an important, yet challenging, objective with potential clinical implications. One challenge has been to parse the role of myocardial substrate utilization intrinsic to the molecular regulation of oxidative enzymes given the confounding impact of whole body and extracellular substrate availability and changes in cellular and mitochondrial uptake. Here, we set out to mimic HFpEF by leveraging a transgenic mouse line expressing PDK4 exclusively in cardiomyocytes, blocking conversion of pyruvate to acetyl-CoA that can be metabolized within the TCA cycle. As such, we sought to elucidate metabolic events in the HFpEF heart normally associated with a high-fat diet, without systemic metabolic alterations or those occurring in other, non-cardiovascular organ systems.

FAO in mitochondria accounts for 60% of myocardial energy production^24^. Under conditions of hypertrophic stress, increases in glucose utilization result in a decrease in fractional oxidation of FFA. We observe a 25% reduction in fractional oxidation of FFA after TAC (**Fig 5A**). However, an examination of *absolute* rates of substrate oxidation shows that the amount of FAO is only slightly decreased (**Fig 5C**). In fact, most of the “switch” is the result of the doubling of glucose/pyruvate oxidation, fueling the increased cardiac demand (**Fig 5C**). Surprisingly, in the absence of glucose/pyruvate oxidation as a substrate option, PDK4Tg TAC hearts still manifest a decrease in fractional oxidation of FFA (**Fig 5A**) driven in this case by an increase in the use of endogenous unlabeled substrates. Absolute levels of FAO do not increase either under these conditions and instead manifest a slight decrease similar to that observed in WT TAC hearts (**Fig 5C**). Additionally, total TCA flux is also decreased in the PDK4Tg hearts.

This unanticipated result, that under conditions of modestly increased cardiac workload wherein glucose oxidation is inhibited, the absolute level of FAO does not increase but paradoxically goes down, mimics a recent observation in human HFpEF hearts^15^. In that study, changes in metabolites from human HFpEF and HFrEF endomyocardial tissue and plasma were examined^15^. Surprisingly, the results showed a decrease in fatty acid metabolites in the hearts of HFpEF patient hearts, as compared to HFrEF, even though glucose oxidation is inhibited in these hearts. These changes were not detectable in the plasma^15^. Furthermore, TCA metabolites on the whole were lower in HFpEF hearts, suggesting substantial disruption of substrate utilization^15^. These data suggest that PDK4Tg hearts are metabolically similar to HFpEF hearts.

PDK4Tg mice provide an opportunity to examine changes in substrate utilization independent of lipid overload. Previous studies have shown that diabetic human hearts, as well as hearts from animals fed a high-fat diet long term, develop insulin resistance which decreases glucose oxidation by not only upregulating PDK4 expression but also decreasing uptake of glucose into the cell. This leads to reliance on FAO that has previously been described as an increase in FAO. Our findings, however, demonstrate clearly that FAO reliance does not equal FAO upregulation. We did not detect signs of lipotoxicity, such as increased ROS or toxic lipid intermediate accumulation, supporting that lipotoxic changes derive from increased FFA availability rather than from increased FFA oxidation.

Carnitine is required for the translocation of long-chain fatty acids across mitochondrial membranes^25^, and previous studies have implicated diminished levels of carnitine in hypertrophied myocardium to the inhibition of FAO after TAC^26^ ^27^. In our hands, the maximal rate of enzyme activity of short-, medium-, and long-chain FFA acyl dehydrogenases (ACAD) from isolated mitochondria of PDK4Tg mice were similar to those in WT hearts (data not shown). This spectrometric experiment is conducted with excess quantities of carnitine and substrates so the protocol bypasses questions of substrate availability to evaluate the regulation of the enzyme activity within mitochondria (assuming no disruption in tertiary configurations and post-translational modifications during isolation). We observed that protein levels of ACAD for every FFA decreased following TAC, and all except long-chain ACAD decreased in the PDK4Tg. Interestingly, mRNA levels of ACADm were the most significantly affected by TAC, in direct contrast to the carnitine-depletion hypothesis. In addition, mass spectrometric measurement of carnitine in LV tissues of WT and PDK4Tg animals post-TAC were largely unchanged with a VIP (variable importance in projection) score of 1.04 and a VIP threshold of 1 for statistical significance. When WT sham animals were used as the control group, carnitine in WT TAC animals were 0.94-fold of control, i.e. only 6% decrease with TAC. PDK4Tg animals harbored higher carnitine amounts: PDK4Tg Sham at 1.43-fold control and PDK4Tg TAC at 1.2-fold control. Therefore, any carnitine decrease due to TAC would have been overcome by its upregulation due to PDK4 overexpression. In aggregate, these findings suggest that changes in carnitine abundance are unlikely contributors to the phenotype observed in our experiments.

Our findings support the notion that PDK4Tg hearts are metabolically similar to HFpEF hearts. That said, the PDK4Tg mice are uniquely susceptible to moderate increases in afterload demand. Indeed, mice fed a HFD and treated with L-NAME develop a HFpEF, and not HFrEF, phenotype^16^. The basis for this is unclear; however, it may be due to long-term adaptation as these animals are metabolically inflexible from birth and HFD feeding of the PDK4Tg mice along with L-NAME afforded no protection (results not shown). The PDK4Tg hearts are reliant on FAO for mitochondrial energy generation and our data suggest that this is sufficient for baseline cardiac demand even without pyruvate utilization. Proteomics data suggest a decrease in the steady state level of a number of FAO enzymes, and TCA flux analysis shows a decrease in the TCA cycle. In fact, the absolute level of FAO in these hearts is comparable to WT hearts at baseline with the decrease in TCA flux arising from lack of glucose/pyruvate oxidation. While sufficient at baseline, this level of FAO is inadequate to support contractile function in the setting of mildly elevated afterload, either from L-NAME or TAC surgery^28^. Surprisingly, both WT and PDK4Tg mice manifest a decrease in gene expression and steady state levels of FAO proteins post-TAC. The PDK4Tg animals displayed minor increases in glycolysis and pyruvate utilization and larger changes in oxidation of endogenous substrates in response to increased demand, suggesting independent regulatory mechanisms of fat and glucose oxidation. This is in addition to the well-established substrate homeostasis governed by the Randle cycle, in which increased use of one substrate inhibits use of the other^29^.

In our hands, suppression of pyruvate metabolism did not impact hypertrophic growth of the myocardium triggered by increased afterload elicited by L-NAME or by TAC surgery, although PDK4Tg mice did progress rapidly to systolic dysfunction in response to either stimulus. Both of these observations are consistent with findings in the setting of PDK4 transgenesis coupled with over-expression of constitutively active calcineurin A (CnA) in which increased cardiac fibrosis and necrosis were observed without exacerbated cardiomyocyte hypertrophy^6^. These findings highlight the independent regulation of hypertrophic and metabolic remodeling in response to increased afterload.

### Study limitations

In an effort to glean insights into myocardial metabolism in HFpEF and to study the effects of metabolic inflexibility on cardiac function, we employed a strategy of transgenic over-expression of PDK4 from birth; yet, this increase in PDK4 activity exceeds that observed in HFpEF. However, our data nonetheless suggest specificity of substrate phosphorylation. Additionally, whereas our ex vivo NMR analyses evaluate flux of metabolites and provide more information than a one-time snapshot, these results may not completely reflect *in vivo* conditions. Furthermore, the number of substrates that can be labeled simultaneously is limited. Therefore, we were able to test only one species of lipid, and our results cannot differentiate between pyruvate and lactate. The latter point is unlikely to be of great significance, however, as these compounds are in equilibrium and used by cardiomyocytes to similar degrees.

### Summary and perspective

The switch between reliance on FAO to glucose oxidation in response to increased cardiac load has long been described as a “preference” and suggested to be the result of changes in oxygen availability. Our data suggest that the inability to upregulate FAO in the face of increased energy demand may underlie the switch to glucose oxidation. Deciphering mechanisms controlling this shift and limiting the pivot to FAO may provide insight into the metabolic changes that occur in the hearts of HFpEF patients, that likewise lack the option to upregulate glucose oxidation in the face of increased demand.

## Supporting information

Piristine data supplement

## ACKNOWLEDGMENTS

The authors acknowledge the assistance of the UT Southwestern Electron Microscopy Core, as well as Dr. Steven A. Kliewer for providing PDK4 transgenic mice. This work was supported by grants from the NIH: HL-128215 (JAH), HL-147933 (JAH), HL-155765 (JAH, TGG), HL-164586 (JAH, TGG), S10RR023729 (JAH) and 1S10OD021685-01A1 (Katherine Luby-Phelps).

