## Supplementary material for "Afterload-induced Decreases in Fatty Acid Oxidation Develop Independently of Increased Glucose Utilization": Piristine data supplement

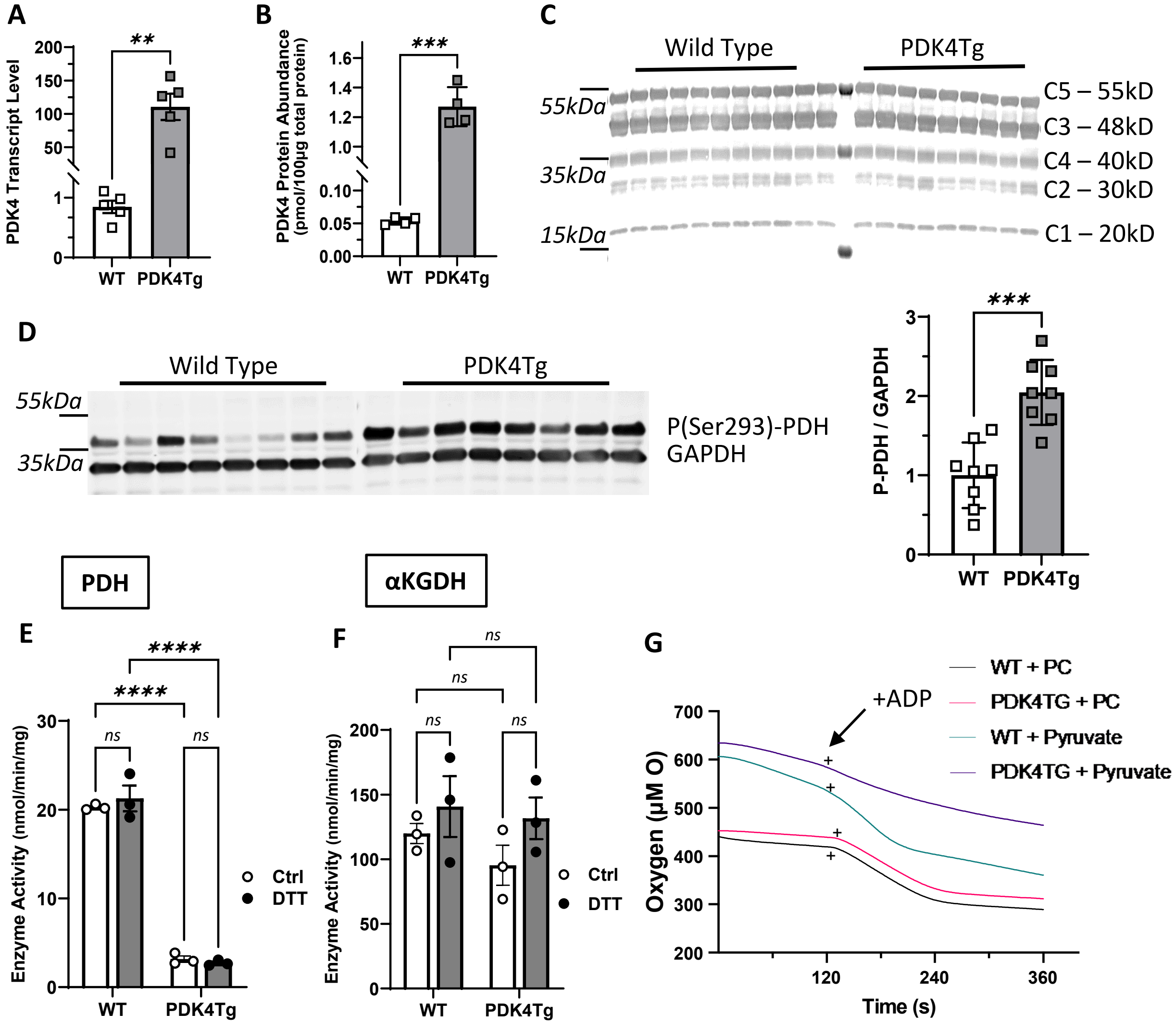


Supplemental Figure 1

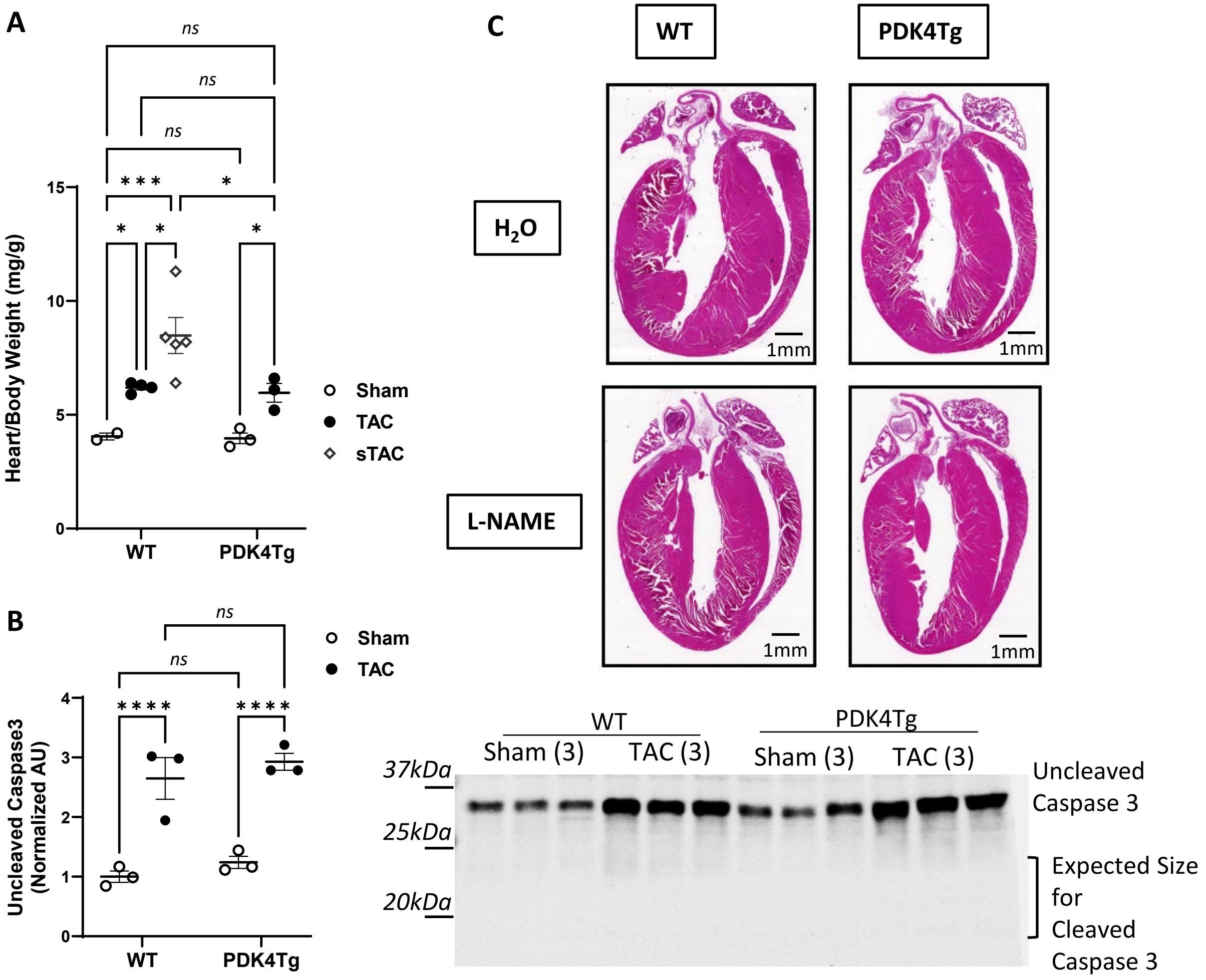


Supplemental Figure 2

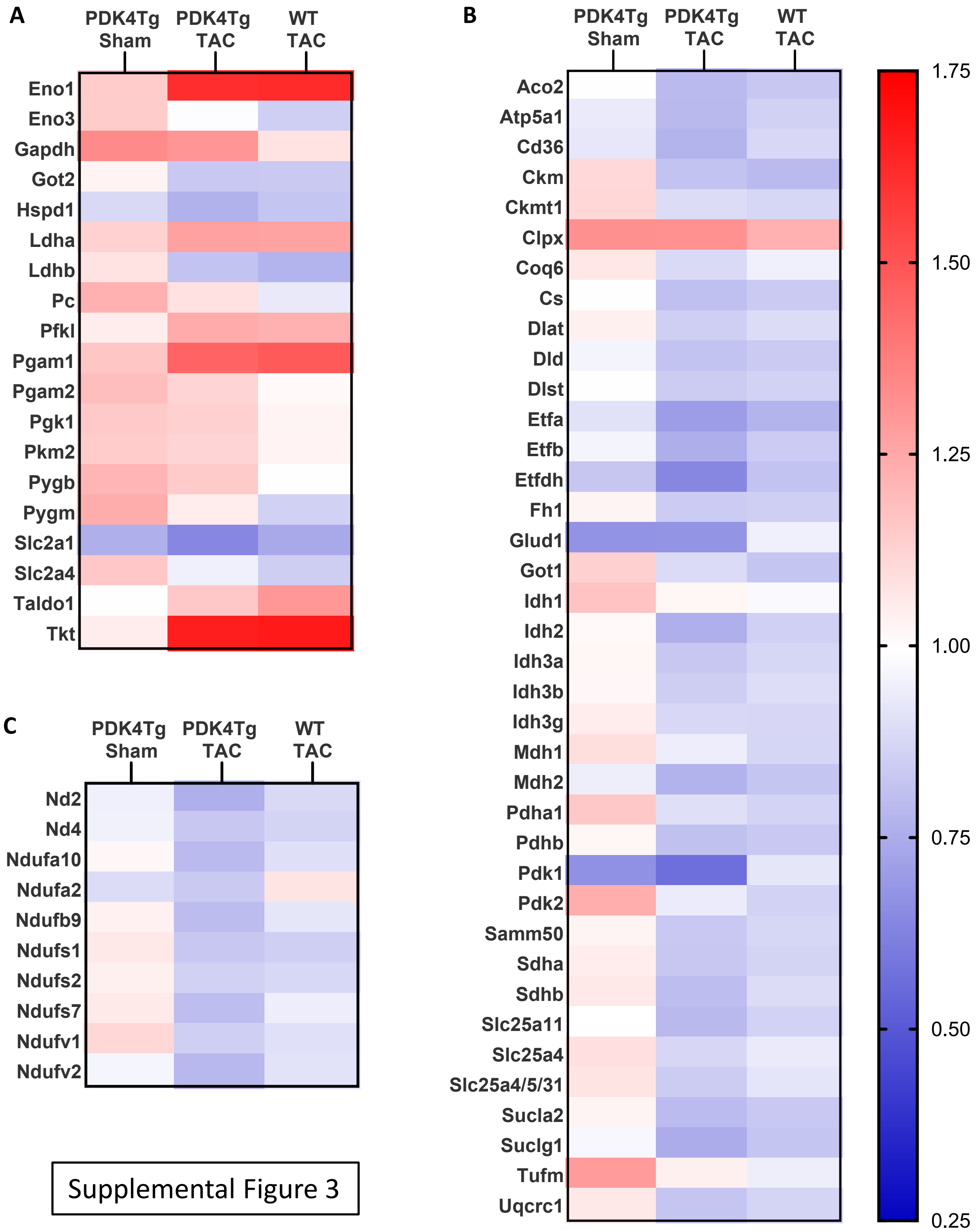


Supplemental Figure 3

**SUPPLEMENTAL FIGURE LEGENDS**

**Supplemental Figure 1.** Molecular phenotype of PDK4Tg mice

A. RT-qPCR measurement of PDK4 gene transcript levels in the LV of 12-week-old PDK4Tg mice and their WT littermates. Expressed as fold change over WT controls after normalizing to mean of control genes Eef1e1 and Rpl4. B. Mass spectrometry quantification of PDK4 protein levels in the LV of 12-week-old PDK4Tg mice and their WT littermates, normalized to total protein in each sample. C. Western blot analysis of PDH E1 alpha subunit Ser293 phosphorylation in WT and PDK4Tg mice whole heart homogenates. GAPDH shown for loading control. n=8. D&E. Spectrometry measurement of enzyme activity in subsarcolemmal mitochondria isolated from WT and PDK4Tg mouse hearts, with or without the incubation with DTT before assay. Activity quantified over time per amount of mitochondrial protein loaded as the rate of NADH production at 340nm from $NAD^+$ with addition of substrate. F. Western blot analysis of ETC complexes I through V in non-denatured whole heart homogenates from WT (n=11) and PDK4Tg mice. G. Oxygen consumption rate of mitochondria isolated from WT and PDK4Tg mouse hearts measured by a fluorescence-based oxygen sensor chamber. Mitochondria were provided with excess amounts of either pyruvate or palmitoyl carnitine (PC) before addition of ADP at 2 minutes to initiate state 3 respiration. For all above, asterisks indicate number of zeroes in p values of Tukey post-hoc multiple comparison tests (MCT) in 2-way ANOVA: *= p≤.05, **= p≤.005, ***= p≤.0005, ****= p≤.00005. *ns* = not significant, p>.05.

**Supplemental Figure 2.** Effect of hypertrophic remodeling on PDK4Tg mice tissue health.

A. Quantification of cardiac growth in WT and PDK4Tg mice 3 weeks post-surgery expressed as heart weight normalized to total body weight at the time of sacrifice. TAC: transverse aortic constriction, sTAC: severe TAC, sham: control surgery without banding of aorta. B. Representative Western blot and quantification of caspase 3 protein levels in WT and PDK4Tg mouse LV tissue 1 week after surgery. Graph shows densitometry measurements of uncleaved caspase 3 (35kDa) normalized to total protein signal per lane. Cleaved caspase 3 bands are expected at 17 & 19kDa per supplier (CST Caspase-3 (D3R6Y) Rabbit mAb #14220) but were not observed. For A. & B., asterisks indicate number of zeroes in p values of Tukey MCT in 2-way ANOVA. C. Representative images of H&E-stained sagittal tissue sections from WT and PDK4Tg mice hearts collected after 6 weeks of consuming regular water ('$H_2O$') or water containing 0.5mg/mL L-NAME ('L-NAME').

**Supplemental Figure 3.** Changes in levels of proteins involved in metabolism with PDK4 overexpression and pressure overload.

Heatmap representation of changes in expression of proteins involved in glycolysis, A., TCA cycle, B., and ETC, C. measured by mass spectrometry in WT and PDK4Tg mice 1 week after sham or TAC surgery. Only proteins whose levels changed statistically significantly (p<.05) either due to surgery or PDK4 gene overexpression by 2-way ANOVA are visualized. Mean values of each protein were calculated for each group (n=4) before being divided by mean value for that protein in WT sham group. Color corresponds to fold change over mean value of WT sham mice for that protein. Red = expressed more, blue = expressed less compared to WT sham. Range of color scale applicable to all 3 images and determined by maximum and minimum value over all 3 pathways. (Note that PDK4 protein levels significantly changed but are not depicted to limit gradient range. PDK4 protein data from this experiment can be found in Fig S1B.)
